# Assessment of spring wheat responses to late season heat stress under Mediterranean conditions

**DOI:** 10.1101/2025.07.29.667146

**Authors:** Shaimaa Mahmoud Awad-Allah, Samer Mohamed Amer, Mahmoud Hassan Abdel-Moneim

## Abstract

In Mediterranean climatic regions, wheat yield is significantly influenced by sowing date and cultivar selection, particularly where production could be limited by heat waves during critical growth stages. This investigation was undertaken to determine the effect of shifting sowing date on a cohort of six high yielding spring wheat cultivars (*Triticum aestivum* L.), by delaying sowing date 15 and 30 days in a two-year field trial, aiming to highlight agronomic traits contributing to sustained yield potential under heat stress induced by late sowing, with a focus on identifying tolerant cultivars.

Heat stress during anthesis reduced grain set and yield by up to 42%, mainly via reduced grain number per spike and thousand grain weight Late sowing shifted the timing of heat stress, resulting in floral abortion and significantly reduced number of grains/spike. More importantly, some cultivars showed up to 10% significant reduction in thousand grain weight (TGW) following delaying sowing date by 30 days, together with the TGW strong correlation with yield especially at late sowing (r=0.74), imposing the yield penalty in the heat stressed late sowing conditions. Among the cohort of tested cultivars, Sids12 and Misr1 cultivars showed minimal yield reduction (10-14%) due to 30 days delayed sowing in the second season highlighted with hot spring.

On the contrary, harvest index of sowing delayed by 30 days was significantly higher compared to both 15 days late sowing and the optimum sowing date, suggesting that decline in biological yield exceeded that of grain yield.

Upon shifting sowing date, the strength of traits association is altered, grain yield associations showed the strongest positive correlation in the 30 days late sowing with plant height (r=0.86), in the 15 days late sowing with number of total tillers/m^2^ (r=0.63) and in optimum sowing date with biological yield (r= 0.62), suggesting that grain yield enhancement through selected traits with direct or indirect contributions should be evaluated independently across different growing conditions.

## 2 Introduction

Wheat (*Triticum aestivum* L.) is globally feeding 2.5 billion people in more than 85 countries and a source of at least 20% calories in the human diet in these countries [1], Moreover, it accounts for 40% of the protein and 33% of the calories in the Egyptian diet [2].

Wheat yield is a polygenic quantitative trait influenced by multiple components, *viz*., number of spiked tillers, number of grains per spike, and grain weight. It is also highly sensitive to environmental conditions [3]. One critical agricultural practice for maximizing yield is identifying the optimal sowing date. Delayed sowing can negatively impact yield, its components, and overall wheat growth and development [4]. Specifically, it can disrupt key growth stages like heading time, grain filling rate and maturity[5], leading to reduced number of spikes per plant and number of grains per spike [6], ultimately causing yield penalty. However, the significance of these effects profoundly varies depending on the genotype [7].

Climate change poses significant threats to wheat production. It was reported that global temperatures are rising by 0.18°C per decade, with each degree of increase potentially reducing wheat production by 6% [8], with several empirical and simulated studies reporting 6.5%-27% yield reduction with above optimum temperature increase during various critical growth stages [9-11]. Therefore, a key focus for researchers is enhancing yield and stability while minimizing resource investment. Hütsch et al. [12] found that number of grains per spike was reduced under heat stress by 83% and 75% during grain filling and at maturity, respectively but decreased the individual grain weight only by 23% at maturity and that lead to a significant 36% reduction in harvest index.

Wheat is considered a heat sensitive crop [13] having optimal temperature of about 20°C for above ground growth and development [14]. Late sowing could put wheat under stress, specifically terminal heat stress which adds yield penalty because of reduction in grain yield key components *viz*., tillering potential, number of spikes/unit area, number of grains/spike and grain weight [12]. Heat stress during the terminal growth stage accelerates the assimilation of photosynthates due to early senescence, leading to smaller grains and reduced quality [15].

Number of grains per spike is established during and shortly after anthesis, mainly via the process of grain setting. A process affected by the maximum number of florets initiated at the double-ridge stage, which occurs simultaneously with the onset of tillering, and the proportion of these florets successfully developing into grains after pollination [16]. Consequently, this yield-determining factor is also partly shaped by events that take place during the vegetative growth phase. Heat stress can negatively impact the formation of double ridges on the shoot apex [12], impede fertilization due to pollen sterility [17] and cause the abortion of already fertilized grains because of inadequate assimilates supply.

The rate and duration of grain filling are the main drivers of grain weight. Research investigating the effects of heat stress during this phase showed that while the grain filling rate increases, the duration is significantly shortened. Wheat plants exposed to heat stress reached maturity earlier than those grown under optimal conditions. However, the yield losses caused by the reduced grain filling period were not compensated for by the accelerated filling rate [18-20].

Therefore, this study aimed to (i) quantify the impact of late sowing on yield and yield components, (ii) identify traits most associated with yield under heat stress, and (iii) highlight cultivars tolerant to late-season stress.

## 3 Materials and methods

### 3.1 Plant material

The used germplasm is a collection of six spring wheat cultivars, obtained from Ministry of Agriculture, Egypt, listed in Table 1.

**Table 1.**
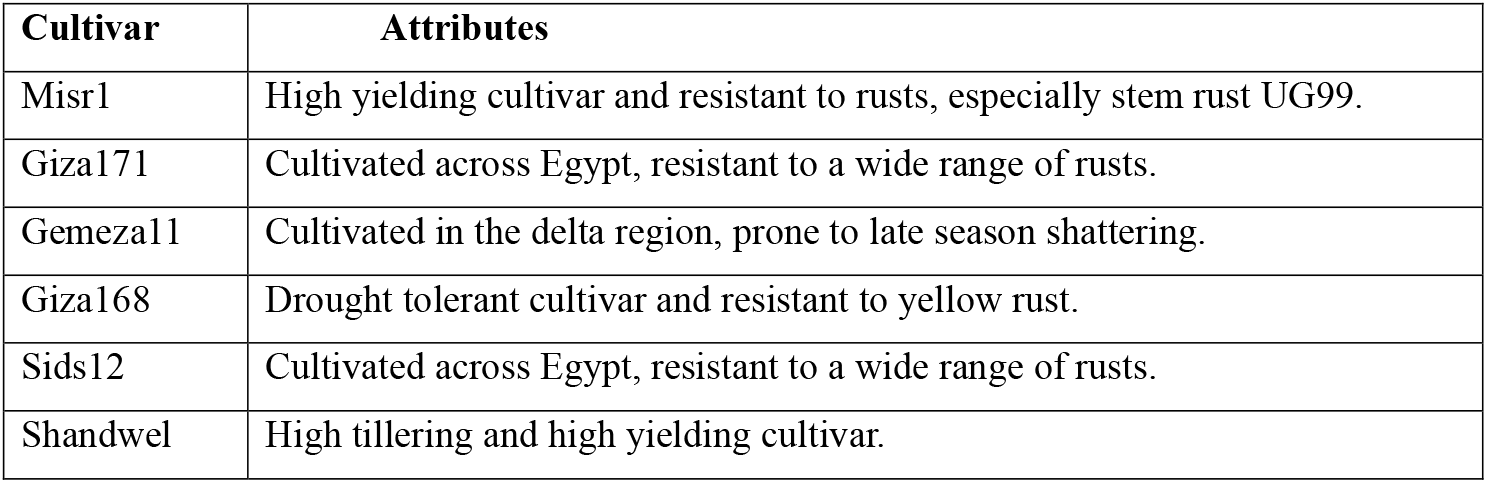
Particulars of Egyptian spring wheat cultivars used in the study.

### 3.2 Field trials

The genotypes panel was evaluated during 2018-2019 and 2019-2020 winter seasons at the experimental station of the Faculty of Agriculture, Alexandria University, Alexandria, Egypt (31°20, N, 30° E), where the soil is sandy loam (54% sand, 30% silt, and 16% clay), electrical conductivity of 1.30 dS m^-1^ and pH 8.05, average and maximum monthly temperature (°C) for the two seasons are illustrated in Figure 1.

**Figure 1.**
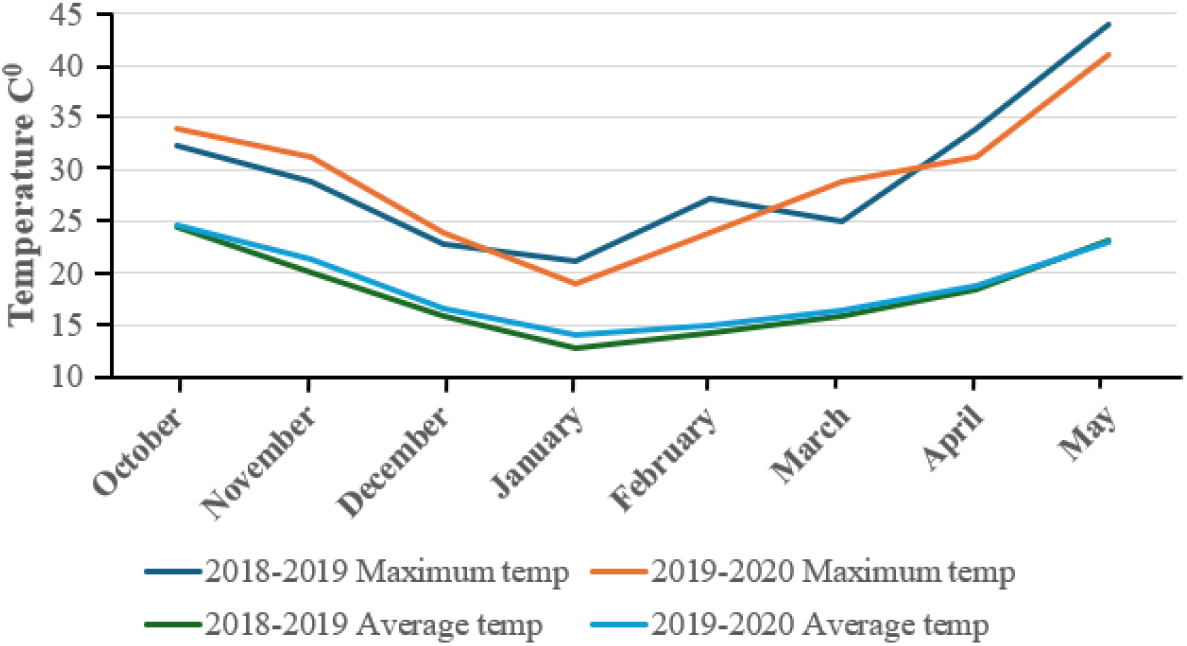
Maximum and average temperature for the two growing seasons.

A randomised complete block design was employed in a split plot, consisting of three blocks (replicates). The main plots were assigned for the three sowing dates, with 15 days interval (15^th^ November, 30^th^ November and 15^th^ December), noting that optimum sowing date under such environment is mid-November, while the sub-plots were dedicated for the cultivars. Seeds were sown in 4 m^2^ (2 x 2m) plots and seeding rate of 57 g/plot (143 kg/ha). Plots were maintained free of weeds and disease with the appropriate herbicides and fungicides and received standard irrigation and fertilization regime (Nitrogen fertilization as 476 kg/ha of urea 46.5% split over three doses, Phosphorus as 595 kg/ha of mono-superphosphate 15% added to the soil during seed bed preparation and Potassium as 119 kg/ha of potassium sulphate added 30 days after sowing.

### 3.3 Phenotyping

Preharvest traits; plant height (P.ht) was measured in cm using the rising plate [21] as the average of two measurements per plot at maturity, number of total tillers/m^2^ (TT) measured as number of tillers in randomly picked 1m^2^ within the plot and number of spiked tillers/m^2^ (ST) measured as number of tillers bearing spikes in the same randomly picked 1m^2^. Post harvest traits; spike length (SL), number of spikelets/spike (SS) and number of grains/spike (GS) measured as the average of ten spikes/plot randomly collected before harvesting, manually measuring the spike length and count number of spikelets/spike then threshed to estimate grain yield, biological yield (BY) by manually harvesting all above ground biomass/plot at maturity and grain yield (GY) quantified by mechanically threshing the harvested biological yield then weighing the grain yield, both estimated as kg/plot then transformed to ton/hectare, thousand grain weight in grams (TGW) measured as the average of two samples/plot collected from the harvested grain yield. Harvest index (HI) was estimated as grain yield/biological yield * 100.

### 3.4 Statistical analysis

R software R 3.3.4 [22] was used to analyse experimental data. Means and standard errors were calculated using ANOVA to identify differences between treatments, R package ‘Agricolae’ [23] was used to perform split plot analysis for traits without homogeneity of error across years, ‘lmerTest’ R package [24] for combined analysis across years for traits with homogeneous error,’ Corrplot’ package [25] for correlation matrix and heat maps and R package ‘ggplot2’ [26] for graphical illustrations.

## 4 Results

The significant levels of measured traits responses under the effects of the two studied factors and their interactions are presented in Table 2 for the traits with heterogeneous experimental error across years, where the main effect of sowing date, cultivars and their interaction varied among the three traits but showed the same pattern in TGW and GS while SS showed opposite pattern over the two years. TGW was not affected by sowing dates in both seasons but varied significantly among cultivars (Table 3), the significant interactions between sowing dates and cultivars (Table 4) shows Giza 168 and Shandwel with increase in TGW upon delaying sowing, while the other cultivars showed varying degrees in TGW decrease (5-10%) with delayed sowing (Figure 2). Number of grains/spike was significantly affected by cultivars X sowing date interaction, in the first season D2:Giza168 was significantly superior with 58.67 grains/spike and D3:Gemeza11 was the lowest with 30.33 grains/spike, while in the second season D3:Giza168 had the highest grains/spike of 101.67 and D1:Sids12 had the lowest with 57.67 grains/spike (Figure 3).

**Table 2.** Levels of significance of thousand grain weight (TGW), number of spikelets/spike (SS) and number of grains/spike (GS) as affected by sowing dates and cultivars in 2018-2019 and 2019-2020.

| Sources of variation | d.f. | TGW |  | SS |  | GS |  |
| --- | --- | --- | --- | --- | --- | --- | --- |
|  |  | 2018-2019 | 2019-2020 | 2018-2019 | 2019-2020 | 2018-2019 | 2019-2020 |
| **Sowing Date (SD)** | 2 | 86.23ns | 26.31ns | 53.32* | 129.43ns | 931.70* | 693.48* |
| **Cultivar (C)** | 5 | 830.51*** | 481.26*** | 94.12*** | 233.35ns | 1216.09*** | 2812.76*** |
| **SD X C** | 10 | 607.79** | 351.50** | 18.04ns | 1127.71*** | 654.07*** | 1328.52* |

ns: non-significant at $p \leq 0.05$ , \* significant at $p \leq 0.05$ , \*\* significant at $p \leq 0.01$ , \*\*\* significant at $p \leq 0.001$ .

**Table 3.** Mean values for thousand grain weight (TGW), number of spikelets/spike (SS) and number of grains/spike (GS) as affected by sowing dates and cultivars in 2018-2019 and 2019- 2020.

| Trait | TGW |  | SS |  | GS |  |
| --- | --- | --- | --- | --- | --- | --- |
|  | 2018-2019 | 2019-2020 | 2018-2019 | 2019-2020 | 2018-2019 | 2019-2020 |
| **Gemeza11** | 73.45 b | 56.37 b | 23.63 a | 42.78 a | 39.66 c | 73.11 bcd |
| **Giza168** | 68.36 d | 52.47 d | 23.65 a | 40.41 ab | 54.33 a | 89.22 a |
| **Giza171** | 68.74 cd | 52.85 cd | 22.04 b | 37.01 b | 47.55 b | 69.44 cd |
| **Misr1** | 78.54 a | 60.25 a | 22.60 ab | 38.68 ab | 42.55 c | 78.89 b |
| **Shandwel** | 67.00 d | 51.45 d | 23.46 a | 41.90 a | 49.66 b | 75.44 bc |
| **Sids12** | 72.55 bc | 55.75 bc | 19.91 c | 42.31 a | 47.66 b | 67.11 d |
| **LSD 0.05** | 3.98 | 3.03 | 1.08 | 4.65 | 3.62 | 7.58 |
| **D1** | 70.13 | 53.94 | 21.62 b | 42.18 | 50.28 a | 73.94 b |
| **D2** | 73.15 | 55.01 | 22.11 b | 40.97 | 49.39 a | 80.50 a |
| **D3** | 71.06 | 53.83 | 23.92 a | 38.45 | 41.05 b | 72.17 b |
| **LSD 0.05** | ns | ns | 1.17 | ns | 6.33 | 4.46 |

Mean values followed by the same letter (s) are not significantly different.

**Table 4.** Mean values for thousand grain weight (TGW), number of spikelets/spike (SS) and number of grains/spike (GS) as affected by sowing dates and cultivars in 2018-2019 and 2019- 2020.

|  | TGW |  | SS |  | GS |  |
| --- | --- | --- | --- | --- | --- | --- |
| Sowing date and cultivar combinations | 2018-2019 | 2019-2020 | 2018-2019 | 2019-2020 | 2018-2019 | 2019-2020 |
| **D1:Gemeza11** | 71.07 cde | 54.67 cdefg | 21.63 gh | 45.97 abc | 44.00 fghij | 72.67 cdef |
| **D1:Giza168** | 58.93 f | 45.33 h | 23.53 bcdef | 34.30 fg | 54.33 abc | 83.00 abc |
| **D1:Giza171** | 71.07 cde | 54.67 cdefg | 20.80 hi | 35.30 efg | 50.33 cde | 74.67 cde |
| **D1:Misr1** | 80.60 a | 62.00 a | 22.07 efgh | 44.87 abc | 46.00 defghi | 78bcde |
| **D1:Shandwel** | 65.87 e | 50.67 fg | 22.73 cdefg | 47.43 ab | 57.67 ab | 77.67 bcde |
| **D1:Sids12** | 73.23 bcd | 56.33 bcde | 18.93 i | 45.20 abc | 49.33 cdefg | 57.67 g |
| **D2:Gemeza11** | 78.47 ab | 59.00 abc | 23.67 bcde | 39.10 cdefg | 44.67 efghij | 77.33 bcde |
| **D2:Giza168** | 70.93 cde | 53.33 defg | 22.33 defgh | 40.63 bcdef | 58.67 a | 90.00 ab |
| **D2:Giza171** | 68.77 cde | 51.67 efg | 22.73 cdefg | 35.87 defg | 43.67 ghij | 68.33 efg |
| **D2:Misr1** | 80.69 a | 60.67 ab | 21.67 fgh | 36.63 defg | 43.00 hij | 82.33 abcd |
| **D2:Shandwel** | 66.06 e | 49.67 gh | 23.13 cdefg | 43.73 abcd | 54.67 abc | 82.00 abcd |
| **D2:Sids12** | 74.04 abc | 55.67 bcdef | 19.13 i | 49.53 a | 51.67 bcd | 83.00 abc |
| **D3:Gemeza11** | 70.84 cde | 53.67 defg | 25.60 a | 43.27 abcde | 30.33 l | 69.33 defg |
| **D3:Giza168** | 75.24 abc | 57.00 abcd | 25.10 ab | 46.30 abc | 50.00 cdef | 94.67 a |
| **D3:Giza171** | 66.44 de | 50.33 gh | 22.60 defgh | 39.87 bcdefg | 48.67 cdefgh | 65.33 efg |
| **D3:Misr1** | 74.36 abc | 56.33 bcde | 24.07 abcd | 34.53 fg | 38.67 jk | 76.33 cde |
| **D3:Shandwel** | 69.08 cde | 52.33 defg | 24.53 abc | 34.53 fg | 36.67 k | 66.67 efg |
| **D3:Sids12** | 70.40 cde | 53.33 defg | 21.67 fgh | 32.20 g | 42.00 ijk | 60.67 fg |
| **L.S.D** | 6.89 | 5.24 | 1.87 | 8.05 | 6.26 | 13.13 |

Mean values followed by the same letter (s) are not significantly different.

**Figure 2.**
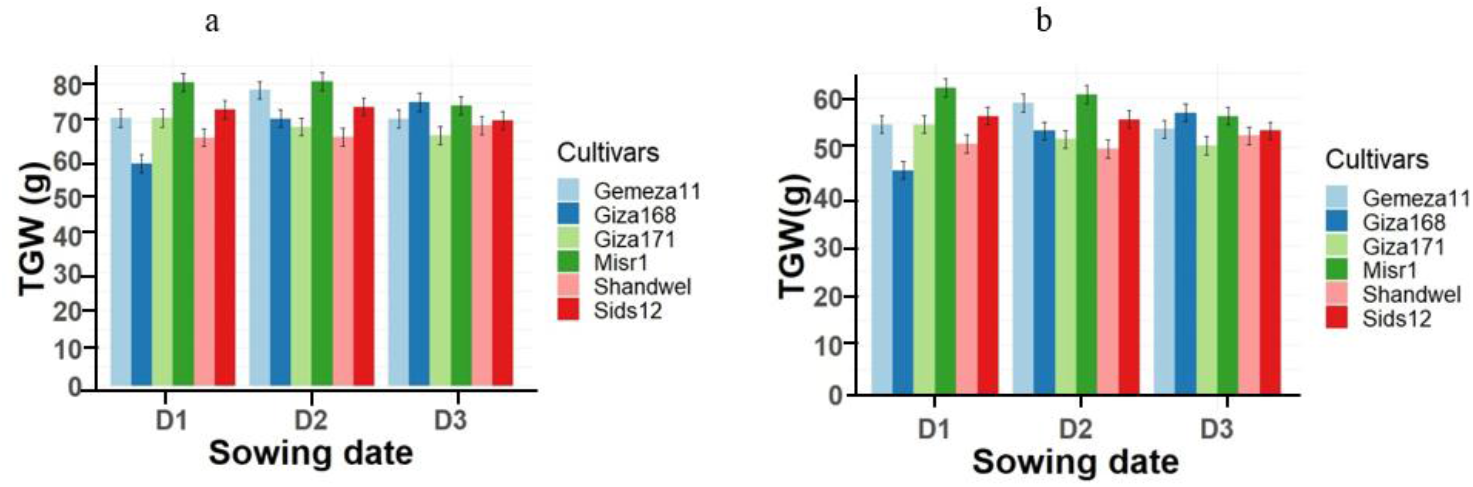
Variations in thousand grain weight (TGW) as affected by the significant interaction between sowing dates and cultivars during (a) 2018-2019 and (b) 2019-2020.

**Figure 3.**
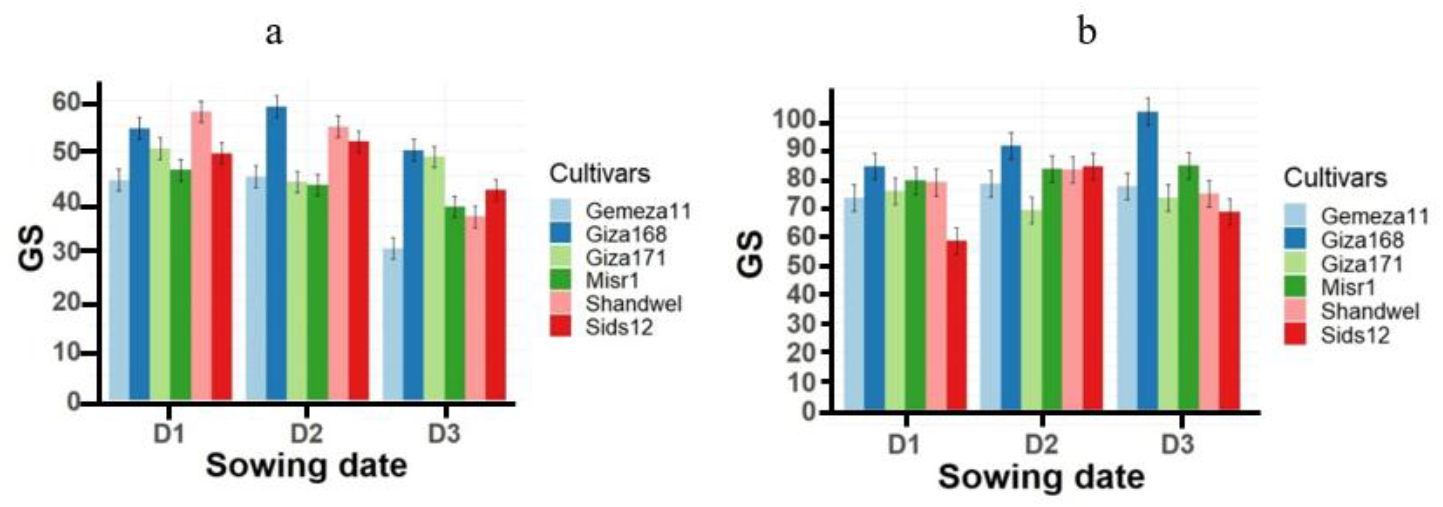
Variations in number of grains/spike (GS) as affected by the significant interaction between sowing dates and cultivars during (a) 2018-2019 and (b) 2019-2020.

In contrast to TGW, number of spikelets/spike in the first season significantly increased by delaying sowing date, as D2 and D3 yielded more SS than D1 by 2.3% and 10.6%, respectively, in the second season cultivars X sowing date interaction was significant, with delaying sowing Giza168 and Giza 171 SS increased, Misr1 and Shandwel SS decreased while Gemezaa11 and Sids12 SS peaked only in D2 (Figure 4).

**Figure 4.**
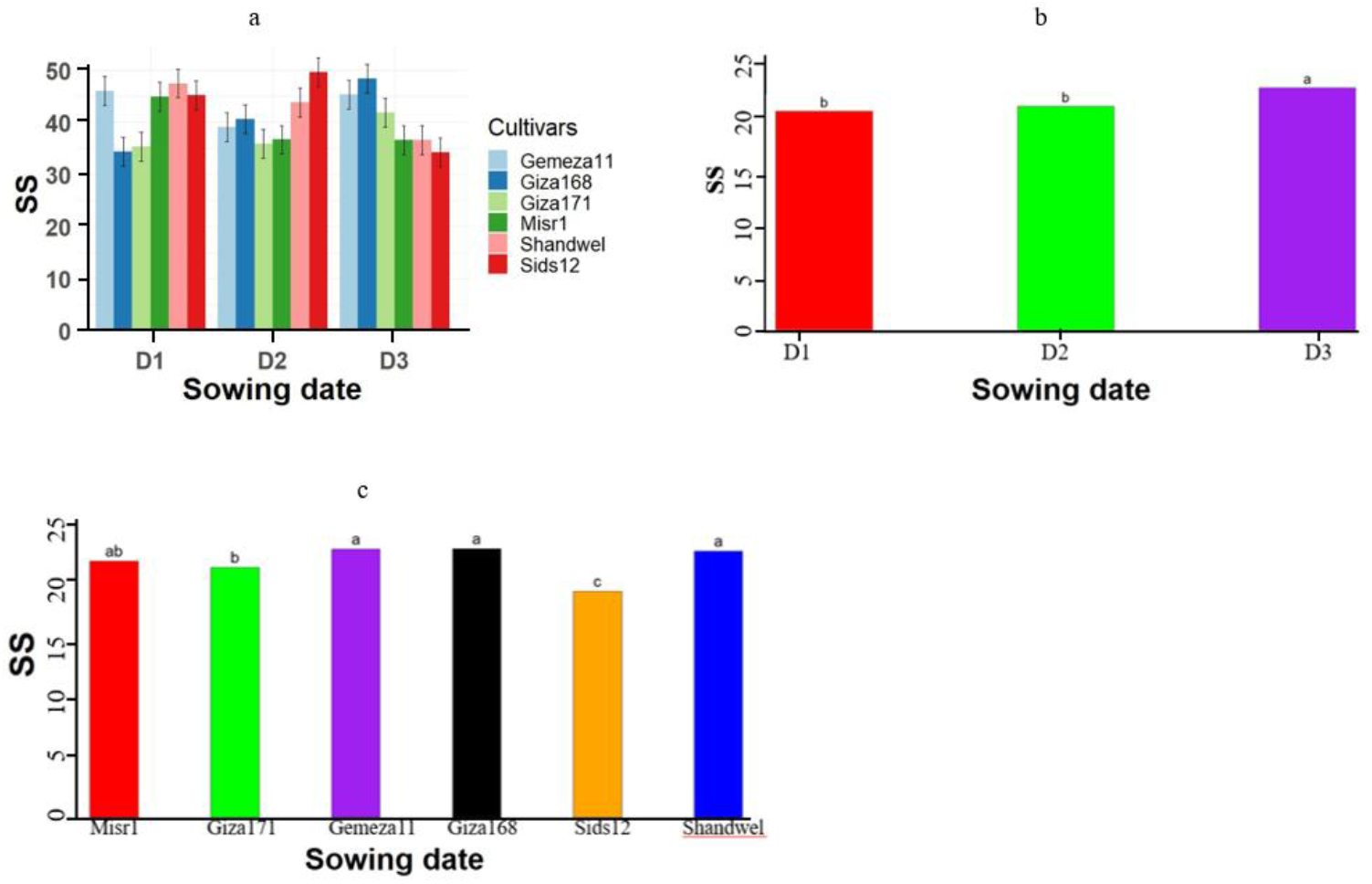
Variations in number of spikelets/spike (SS) as affected by the significant interaction between sowing dates and cultivars during 2018-2019 and by significant main effect of sowing dates and cultivars during 2019-2020.

Table 5 shows that year X cultivar effect was significant for all traits, the three way and SD X C were significant for all traits except plant height, while Year X SD was significant for plant height, biological and grain yield only. The main effect of cultivars was not significant for biological yield, while the main effect of sowing date was not significant for plant height and the year main effect was not significant for number of total tillers and spiked tillers/m^2^.

**Table 5.** Levels of significance of plant height (P.ht), biological yield (BY), grain yield (GY), harvest index (HI), number of total tillers/m^2^ (TT), number of spiked tillers/m^2^ (ST) and spike length (SL) as affected by year, sowing dates and cultivars in 2018-2019 and 2019-2020.

| Sources of variation | D.f | P.ht | BY | GY | HI | TT | ST | SL |
| --- | --- | --- | --- | --- | --- | --- | --- | --- |
| year | 1 | 1891.70* | 9032.70** | 131.71** | 124.38* | 52163ns | 46927ns | 99.57** |
| Sowing date(SD) | 2 | 215.40ns | 1039.70*** | 8.27** | 166.35* | 50398*** | 37041** | 21.44* |
| Cultivar (C) | 5 | 341.80** | 85.70ns | 9.68*** | 272.55*** | 77771*** | 72733*** | 41.91*** |
| Year X SD | 2 | 3373.60*** | 591.70*** | 12.35** | 87.26ns | 312ns | 378ns | 8.89ns |
| Year X C | 5 | 421.00*** | 330.00*** | 13.55*** | 258.16*** | 6554* | 7339* | 24.00* |
| SD X C | 10 | 98.10ns | 760.60*** | 23.29*** | 531.77*** | 22153*** | 29960*** | 37.21* |
| Year X SD X C | 10 | 212.80ns | 396.10*** | 55.45*** | 697.35*** | 10998* | 11810* | 44.30** |

ns: non-significant at $p \leq 0.05$ , \* significant at $p \leq 0.05$ , \*\* significant at $p \leq 0.01$ , \*\*\* significant at $p \leq 0.001$ .

Significance of plant height with Year X Cultivar interaction is mainly due to the stability of Sids12 height across the two seasons while all other cultivars hight was reduced significantly in the second season (Figure 5b). Sowing date effect varied significantly between the two years for D3 (decreased in the second year by 20.5%) while D1 and D2 effect was stable across years (Figure 5a).

**Figure 5.**
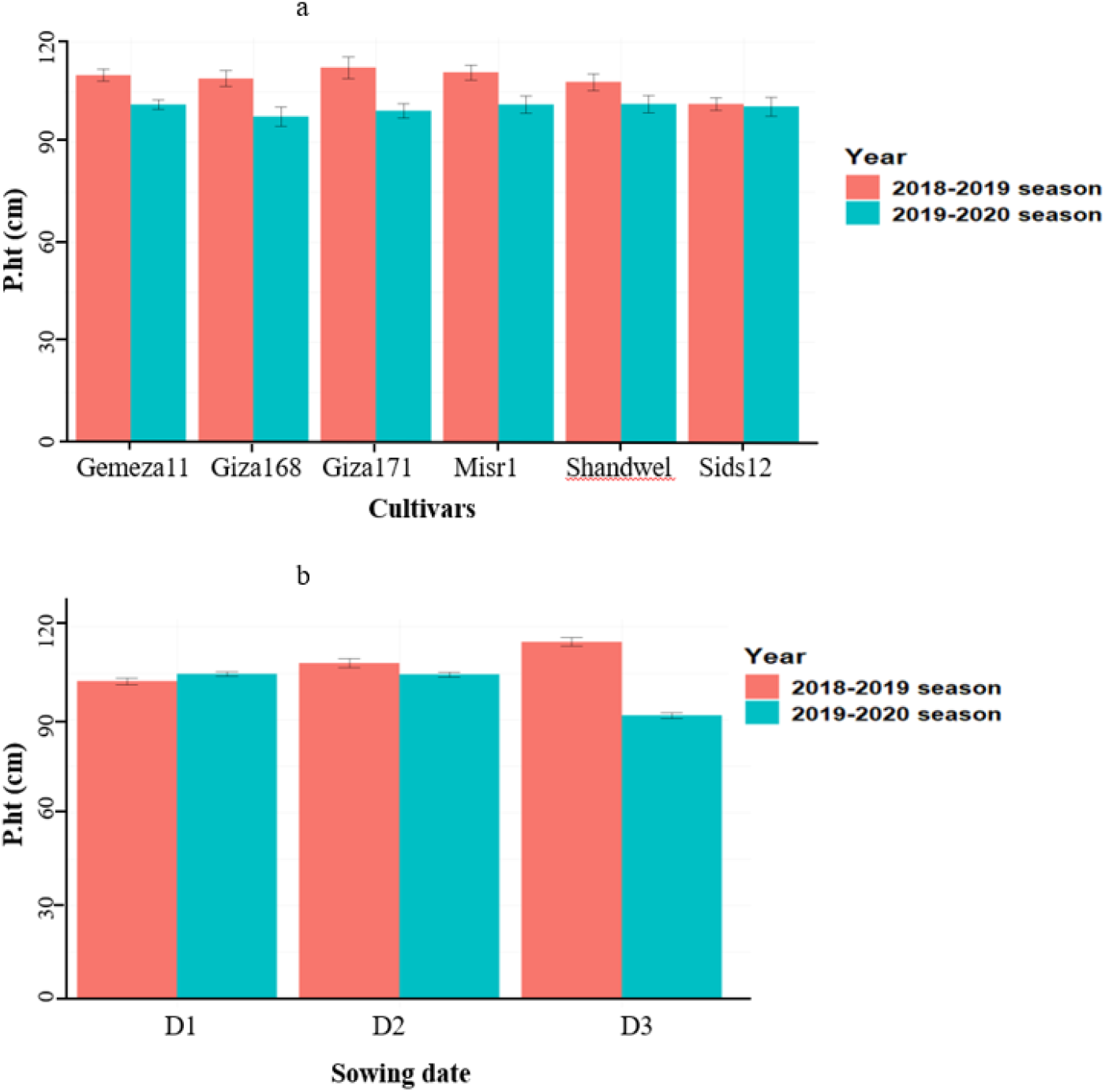
Variations in plant height (P.ht) as affected by the significant effect of sowing dates (a) and interaction between year and cultivars during 2018-2019 and 2019-2020.

The effect of sowing date X cultivar on biological yield varied significantly across years (Figure 7a), highlighting D2:Giza171 in the first year as the highest (42.11 t/ha) and D3:Giza168 in the second year as the lowest (5.30 t/ha), indicating a strong seasonal impact. Notably, all sowing date X cultivar combinations were significantly reduced for biological yield in the second season (except for D2:Shandwel) with various degrees ranging from 28.5% decrease in D1: Giza168 up to 85 % in D3: Giza168 (Table 7).

Table 6 shows significant 42% reduction in yield of D3 compared to D1 and D2 of the second season. The significant Year X SD X C interaction in grain yield showed the superiority of D2:Misr1in the first year (6.85 t/ha) (Table 7) and D3:Giza168 in the second year (same as BY) as the lowest (0.80 t/ha). In the second year, for each sowing date X cultivar combination grain yield was reduced by at least 40% and notably reduced by 83% in D3:Giza168, except for D2:Giza171 which didn’t lose significant grain yield in the second year (Figure 7b). Table 7 illustrates that GY was reduced by delaying sowing date in most cultivars in each season, with few exceptions of increase in D2 or D3 compared to D1 for the same cultivars specially in 2018-2019 season. On the contrary in the second season, all cultivars showed significant yield penalty induced by delaying sowing by 30 days, highlighting Sids12 and Misr1 cultivars with minimal yield reduction of 10 and 14%, respectively.

**Table 6.** Mean values for plant height (P.ht), biological yield (BY), grain yield (GY), harvest index (HI), number of total tillers/m^2^ (TT), number of spiked tillers/m^2^ (ST) and spike length (SL)as affected by sowing dates and cultivars in 2018-2019 and 2019-2020.

| Cultivar/<br>Trait | P.ht |  | BY |  | GY |  | HI |  | TT |  | ST |  | SL |  |
| --- | --- | --- | --- | --- | --- | --- | --- | --- | --- | --- | --- | --- | --- | --- |
|  | 2018-2019 | 2019-2020 | 2018-2019 | 2019-2020 | 2018-2019 | 2019-2020 | 2018-2019 | 2019-2020 | 2018-2019 | 2019-2020 | 2018-2019 | 2019-2020 | 2018-2019 | 2019-2020 |
| Gemeza11 | 110.11 ab | 101.23 cd | 32.02 c | 15.87 de | 4.54 bc | 1.92 f | 14.24 def | 12.61 ef | 243.33 bc | 192.22 gh | 238.00 bc | 183.11 g | 15.10 abc | 15.54ab |
| Giza168 | 109.06 ab | 97.62 d | 35.58 ab | 15.25 e | 5.06 a | 1.88 f | 14.30 def | 13.87 def | 217.11 def | 170.33 hi | 209.11 def | 170.33 gh | 14.25 cd | 15.53 ab |
| Giza171 | 112.31 a | 99.33 cd | 37.73 a | 14.91 e | 4.37 c | 2.28 ef | 12.22 f | 17.11 abc | 199.55 fg | 160.00 i | 194.55 fg | 156.66 h | 12.75 ef | 14.43 bcd |
| Misr1 | 110.84 ab | 101.26 cd | 35.91 ab | 16.28 de | 5.21 a | 2.52 e | 14.96 cde | 16.21 bcd | 254.55 b | 197.55 fg | 243.55 b | 190.22 fg | 12.27 f | 15.23 abc |
| Shandwel | 108.00 b | 101.41 c | 30.08 c | 18.26 d | 4.90 ab | 3.27 d | 17.68 ab | 19.07 a | 300.11 a | 235.11 bcd | 291.77 a | 229.55 bcd | 13.62 de | 15.60 a |
| Sids12 | 101.40 c | 100.64 cd | 34.90 b | 15.89 de | 4.15 c | 3.10 d | 12.09 f | 19.49 a | 230.55 cde | 213.77 ef | 220.55 cde | 207.11 ef | 12.44 f | 15.63 a |
| LSD 0.05 | 3.68 |  | 2.79 |  | 0.47 |  | 2.49 |  | 21.3 |  | 22.02 |  | 1.13 |  |
| D1 | 102.42 c | 104.80 bc | 35.99 a | 20.00 c | 4.89 a | 2.91 b | 14.18 b | 14.87 b | 250.39 a | 204.00 b | 246.56 a | 199.00 b | 12.53 d | 15.23 a |
| D2 | 108.28 b | 104.57 bc | 33.83 ab | 19.76 c | 4.43 a | 2.91 b | 13.83 b | 14.89 b | 209.22 b | 169.33 c | 205.22 b | 166.67 c | 13.39 cd | 15.13 ab |
| D3 | 115.17 a | 91.38 d | 33.30 b | 8.48 d | 4.79 a | 1.67 c | 14.73 b | 19.41 a | 263.00 a | 219.33 b | 247.00 a | 209.33 b | 14.30 bc | 15.62 a |
| LSD 0.05 | 4.3 |  | 2.6 |  | 0.47 |  | 2.81 |  | 22.73 |  | 11.82 |  | 0.92 |  |

Mean values followed by the same letter (s) are not significantly different.

**Table 7.**
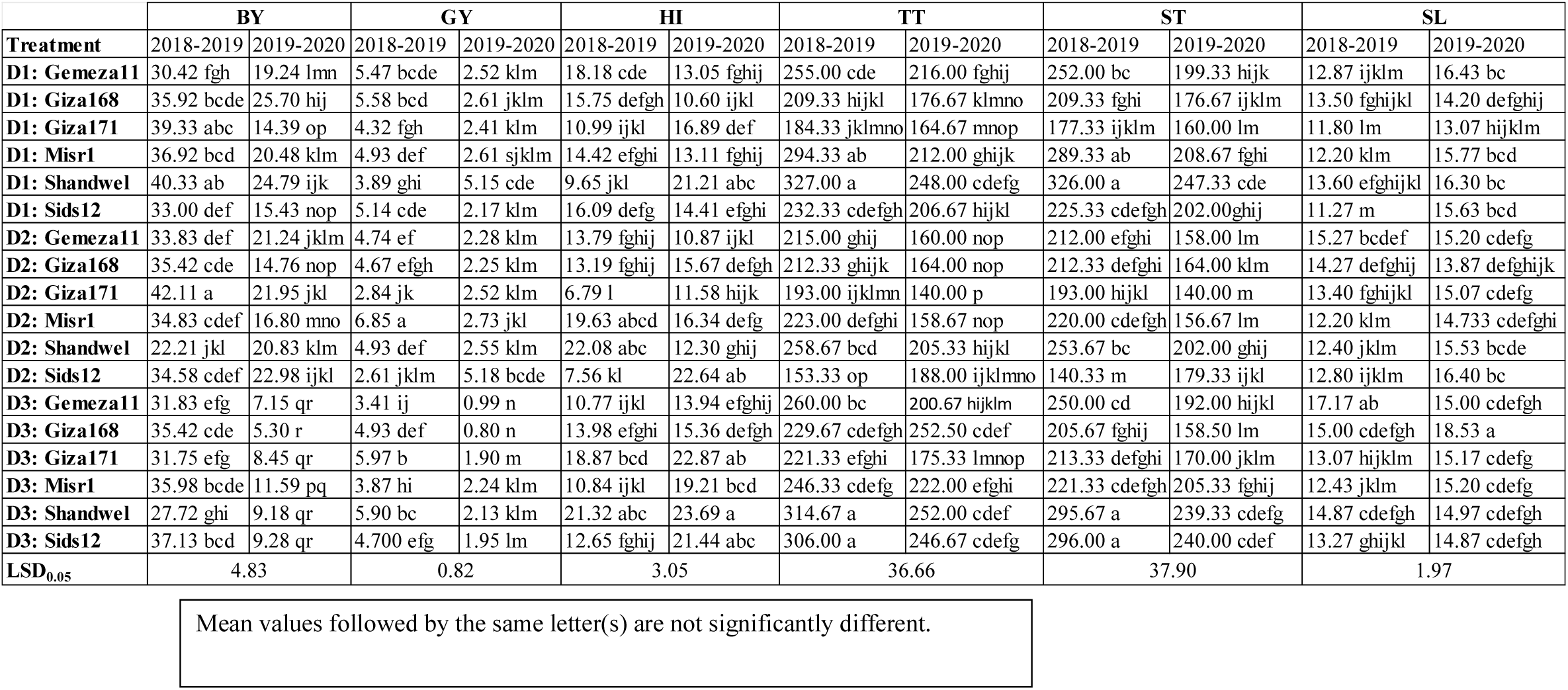
Mean values for plant height (P.ht), biological yield (BY), grain yield (GY), harvest index (HI), number of total tillers/m^2^ (TT), number of spiked tillers/m^2^ (ST) and spike length (SL) as affected by interaction between year, sowing dates and cultivars in 2018-2019 and 2019-2020 seasons.

A different pattern from biological and grain yield was observed in harvest index, although the Year X SD X C interaction was significant, Figure 7b shows that some sowing date X cultivar combinations significantly decreased, others increased, and some didn’t change across years e.g. D3:Shandwel which had the highest harvest index across the two years (Table 7).

Figure 6a illustrates that most sowing date X cultivar combinations were significantly reduced for number of total tillers/m^2^ in the second season, however D2: Sids12 and D3: Giza168 increased insignificantly as shown in Table 7, the significant decrease ranged from 9.8% in D3: Misr1 up to 28.8% in D2:Misr1, noting that the highest decrease for sowing date X cultivar combination between the two years was observed in D2 (20.6%-28.8%).

**Figure 6.**
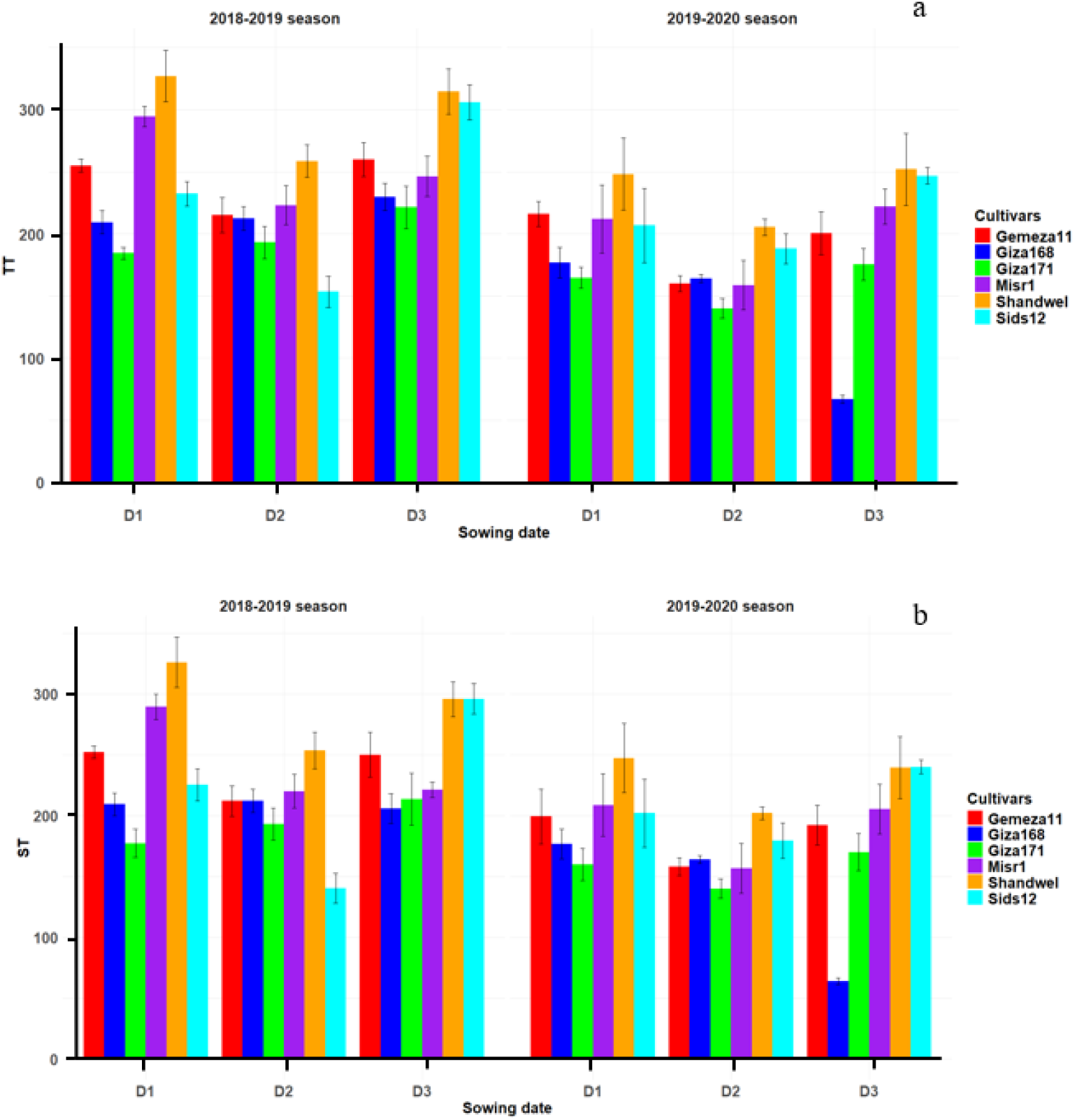
Variations in number total tillers/m^2^ (TT) (a) and spiked tillers/m^2^ (ST) (b) as affected by the significant interaction between sowing dates and cultivars during 2018-2019 and 2019- 2020

A similar pattern for the number of total tillers/m^2^ was found in response of Number of spiked tillers/m^2^ response to the three-way interaction (Figure 6b), with significant decrease in most sowing date X cultivar combinations in the second year ranging from 9.7% in D1:Giza171 up to 28.8% in D2:Misr1, with exceptional one significant increase in the second year for D2:Sids12 combination as 27.8% increase (Table 7).

The significant Year X SD X C interaction in terms of spike length showed D3: Giza168 in the second season with the highest value of 18.53 cm while the shortest spike length of 11.27 cm was found in D1:Sids12 in the first year (Table 7). For most sowing date X cultivar combinations, the spike length significantly increased in the second year from 1.67 cm in D2:Giza171 up to 4.36 cm in D1:Sids12. However, D3:Gemeza11 didn’t follow the same trend and had a significantly decreased spike length in the second year (Figure 7).

**Figure 7.**
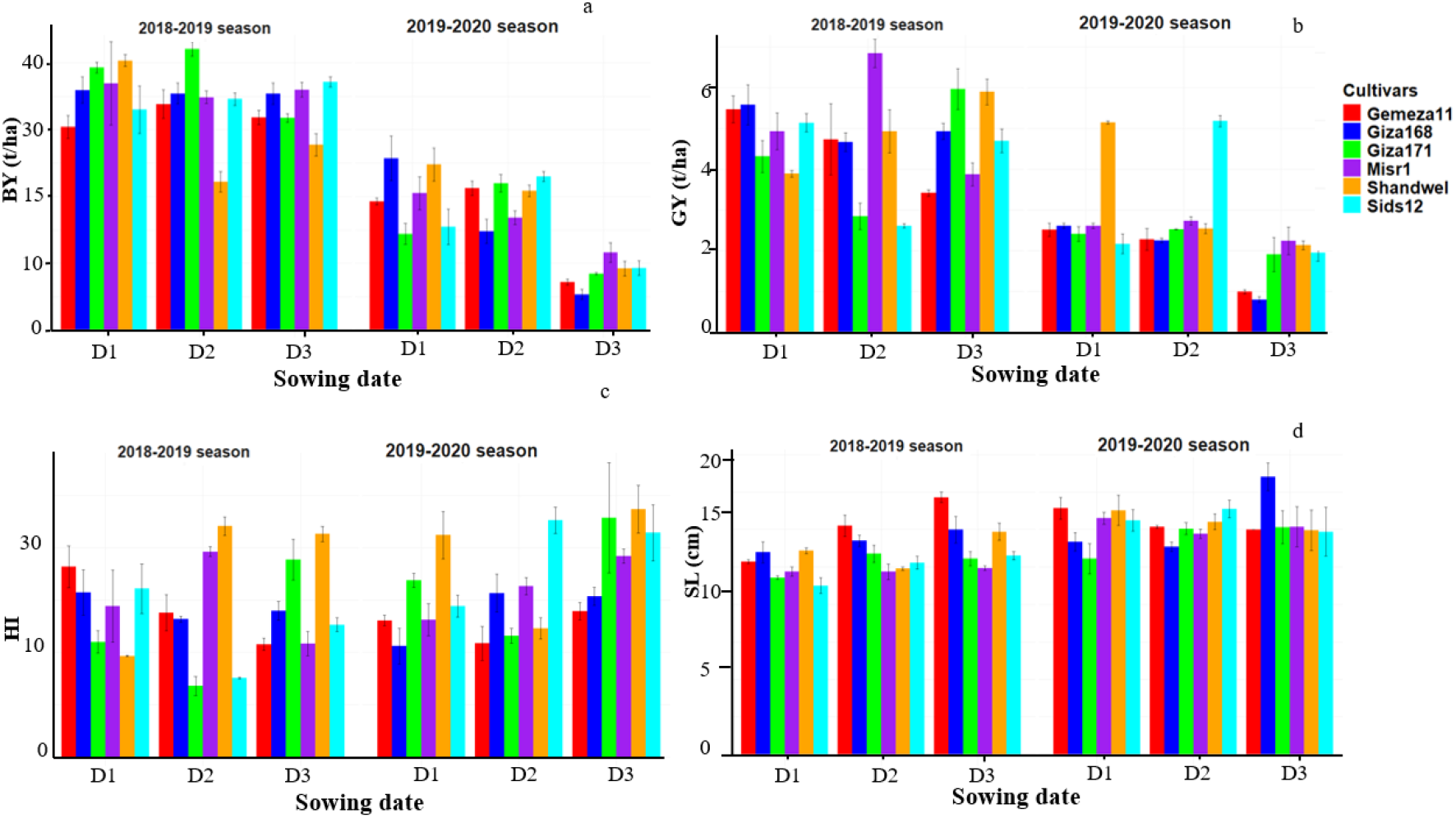
Variations in biological yield (BY) (a), grain yield (GY) (b) harvest index (HI) (c) and (d) spike length (SL) as affected by the significant interaction between sowing dates and cultivars during 2018-2019 and 2019-2020.

Correlations (Pearson’s Correlation Coefficient - PCC) between traits under different sowing dates (Figure 8) were calculated to assess the association among studied traits and the environmental dependences of these associations. Number of total tillers/m^2^ and number of spiked tillers/m^2^ had the highest correlation (r=0.98) over the three sowing dates.

**Figure 8.**
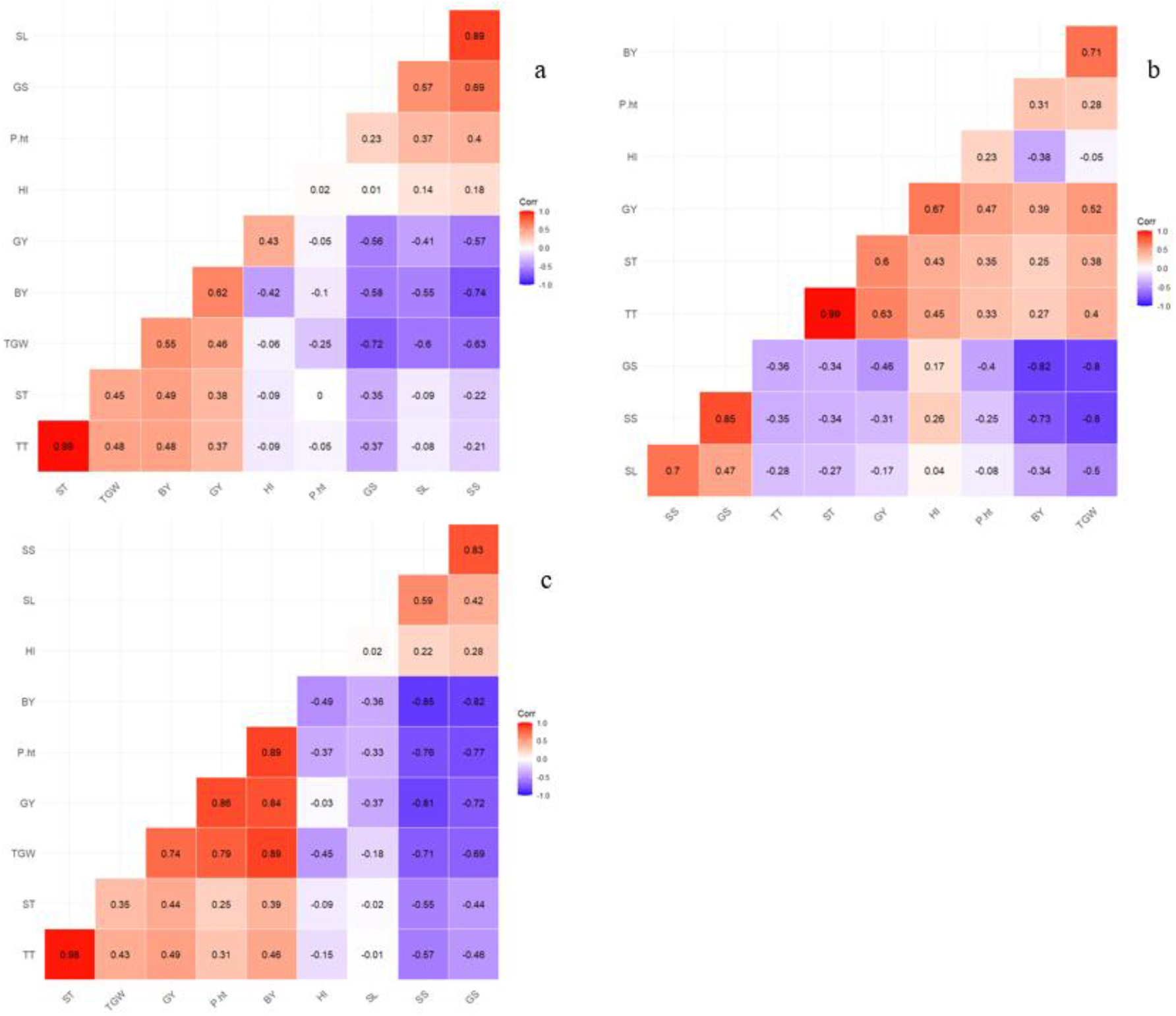
Pearson’s correlation coefficient for the measured traits. (a) D1, (b) D2 and (c) D3.

TGW showed negative association with spike length, number of spikelets/spike and number of grains/spike with various degrees across sowing dates, its strongest (r= -0.8) with number of spikelets/spike and number of grains/spike at D2. On the other hand, number of spikelets /spike and number of grains/spike were both highly correlated and this association peaked (r=0.85) at D2. Figure 8 illustrates the positive correlation between spike length and SS and GS varied among sowing dates, with its highest at D1 (r=0.89 and r= 0.57, respectively). The sowing date effect on association between GS and SS is evidently shown in figure 8, with its lowest association in D1 (r=0.69) and its highest in D2 (r=0.85).

Grain yield associations varied with traits and within sowing dates, noting the strongest positive correlation in D3 with plant height (r=0.86) which fell in D2 (r=0.47) and vanished in D1, while GY positive correlation with ST varied slightly among sowing dates (r=0.38 to r=0.6). Correlations with post-harvest traits showed TGW to have positive association with GY that increased with delaying sowing date, while SS and GS negatively correlated with GY across the three sowing dates (Figure 8).

Harvest index had no significant correlation with GY in D3 but showed significant positive association with GY (r= 0.43) and (r=0.67) in D1 and D2, respectively and showed no significant associations with post-harvest traits across the sowing dates except with TGW in D3 (r=-0.46).

## 5 Discussion

In a Mediterranean type environment the correct choice of sowing date and cultivar are critical determinants of yield, sowing date usually occurs within a sowing window starting with the first significant rainfall in autumn and closing when a sowing date is too late to achieve a reasonable yield [27], with early sowing coinciding soil water availability allowing for longer grain filling duration and maximised leaf area, grain number and size [28]. In this experiment, we investigated the delayed sowing date effect and its consequence heat coinciding with critical growth stages on phenology, yield and yield components of spring wheat.

Although sowing date didn’t affect GY in the first season, we found a significant decline of 42% in the late sowing date D3 in the second season which coincided with above average temperature during late February and March (Figure 1), as early sowing dates increase the interception of solar radiation by the crop, allowing it to accumulate more dry matter and avoid terminal drought/heat at the end of the growing season or shorter harvest window in late season sowing [27, 29], this reduction in dry matter accumulation was reflected in TGW reduction. Moreover, studies considering crop simulations reported earlier sowing dates for late ripening cultivars, allows for a longer growing season and consequently higher yield [30]. In the second year, Sids12 and Misr1 cultivars showed minimal yield penalty as a result for 30 days delayed sowing and its consequent heat stress, which could be mainly explained by their potential to sustain number of fertile spikes/m^2^ which was reduced by only 1.5% in Misr1 and even increased in Sids12, while other cultivar such as Giza186 showed 11% in sustain number of fertile spikes/m^2^.

Number of spikelets/spike in D3 increased significantly in the first season but didn’t change significantly in the second year which can be explained by changes in spike length rather than effect of stress on fertility, as SL showed strong association with SS (Figure 8) and sowing date had the same effect on SL as SS (Tables 3 and 6). Conversely, number of grains/spike decreased significantly in D3, probably due to terminal heat stress during anthesis (Figure 1), which usually takes place at February/early March [31]. GS was reduced in D3 in both seasons by an average of 8.3 grains/spike, possibly due to incomplete fertilization, as heat elevation at anthesis adversely impacts pollens fertility [12, 17]. The number of grains is largely determined by the number of fertile florets, which in turn depends on the dynamics of floret generation and degeneration [32]. Floret degeneration or mortality typically occurs when the stem and spike are growing at their fastest rates [33]. It is believed that the variation in number of grains per spike is linked to changes in the availability of assimilates for spike growth during the period of rapid stem elongation, when floret abortion could occur due to adverse conditions [34]. With limited availability of assimilates, many floret primordia may degenerate or die, leaving less florets reaching the fertile stage at anthesis [33].

TGW didn’t change following sowing date main effect alteration in both years despite the decrease in grains number/spike in the second season, which in normal conditions would have shown the well established trade-off between TGW and GS [35], possibly due to limited source capacity of biomass [36] which appeared as a positively strong correlation (r=0.89) between BY and TGW in Figure 8c. However, heat sensitive genotypes showed 5-10% decrease in TGW upon delaying sowing date adding to the penalty of yield due to delayed sowing.

Biological yield didn’t change in the first season but showed significant 57% diminish in D3 of the second season, matching the pattern of grain yield, these results align with Aslani et al. [6] findings who reported negative impacts of delayed sowing on biomass of wheat. Under stress conditions, the plants adjust to the unfavourable conditions by limiting biomass growth, possibly facilitating increased transpiration while maintaining stable water consumption and water use efficiency [12].

Harvest index was stable in the first season across sowing dates following both BY and GY; Nevertheless, it showed an opposite pattern in the second season as HI in D3 was significantly higher than that of D1 and D2 contradicting several reports which found HI to follow BY and GY decline [11, 12], probably as decline in BY was greater than that of GY.

Plant height showed opposite performance across years, in D3 of the first year it was significantly higher than D1 and D2, an unusual finding reported in one of a two-years trial by Elbashier et al. [37]. In a different study, Sareen et al. [38] found no significant reduction under late sowing/heat stress, conversely, it was significantly lower in D3 of the second year as was most of the measured traits due to heat stress in February/March, agreeing with others who reported shorter plants under late sowing [39, 40].

TT and ST at D3 were equal to that of D1(optimal sowing date) in both years, probably because heat stress induced by late sowing in this experiment didn’t coincide with tillering or booting, while others found it might even increase under late sowing [41, 42]

SL in D3 of the first year slightly surpassed D1, however, in the second year it showed stable performance under the three sowing dates, minimal or no impact of sowing date on spike length was reported by other researchers [41], on the contrary, SL get significantly reduced if the late sowing led to terminal heat stress [43, 44].

As the sowing date changed, the strength of traits association is altered, indicating that grain yield enhancement through prioritize traits with direct or indirect contributions evaluated independently across different growing conditions to account for environmental interactions [29]. This alteration was observed in several cases e.g. SL and GS association which shifted by delaying sowing as r= 0.57, 0.47 and 0.42 in D1, D2 and D3, respectively. Moreover, grain yield associations varied profoundly among sowing dates, noting the strongest positive correlation in D3 with plant height (r=0.86), (r=0.47) in D2 and vanished in D1(r= -0.05) (Figure 8).

The TGW negative association with spike length, number of spikelets/spike and number of grains/spike across sowing dates, this association aligns with the well-established trade-off between TGW, SS and GS [35, 45].

The genotypes showed wide range of responses both within growing seasons and across sowing dates, with significant interactions observed in most of the studied traits. Nevertheless, ‘Giza171’ and ‘Misr1’ demonstrated yield stability in the second year, aligning with their stability in number of grains/spike, highlighting the significance of such trait in sustaining yield potential under heat stress during anthesis, a result reported in investigations on other wheat germplasms [46, 47].

## 6 Conclusion

Heat stress induced by late sowing could significantly impair wheat grain yield, however, the yield components driving the yield penalty significantly vary depending on (1) differential genotypic responses and (2) sowing date-dependent heat stress across growth stages, fluctuating annually. Our results under Mediterranean conditions illustrated that 30 days late sowing could impose heat stress during anthesis, resulting in significant reduction in grain number/spike, thousand grain weight and ultimate grain yield, a consequence that could be alleviated by growing the cultivars with ability of sustaining yield potential under such stress. These results highlight Sids12 and Misr1 cultivars as the most stable cultivars with minimal yield penalty.

## 7 Declarations

### 7.1 Ethics approval and consent to participate

Not applicable.

### 7.2 Consent for publication

Not applicable.

### 7.3 Availability of data and materials

The datasets used and/or analysed during the current study are available from the corresponding author on reasonable request.

### 7.6 Authors’ contributions

The three authors contributed equally to designing, data collection of the experiment, data analysis and writing the manuscript.

## References

1. Yashavanthakumar, K.J., et al., Impact of heat and drought stress on phenological development and yield in bread wheat. Plant Physiology Reports, 2021. 26(2): p. 357–367.

2. Abdalla, A., M. Becker, and T. Stellmacher The Contribution of Agronomic Management to Sustainably Intensify Egypt’s Wheat Production. Agriculture, 2023. 13, DOI: 10.3390/agriculture13050978.

3. Pang, Y., et al., High-Resolution Genome-wide Association Study Identifies Genomic Regions and Candidate Genes for Important Agronomic Traits in Wheat. 2020(1752-9867 (Electronic)).

4. Andarzian, B., et al., Determining optimum sowing date of wheat using CSM-CERES-Wheat model. Journal of the Saudi Society of Agricultural Sciences, 2015. 14(2): p. 189–199.

5. Yasmeen, A., et al., performance of late sown wheat in response to foliar application of moringa oleifera lam. leaf extract. Chilean journal of agricultural research, 2012. 72: p. 92–97.

6. Aslani, F. and M.R. Mehrvar, Responses of Wheat Genotypes as Affected by Different Sowing Dates. Asian Journal of Agricultural Sciences, 2012. 4: p. 72–74.

7. Ahamed, K.U., K. Nahar, and M. Fujita, Sowing date mediated heat stress affects the leaf growth and dry matter partitioning in some spring wheat (Triticum aestivum L.) Cultivars. IIOAB Journal, 2010. 1: p. 8–16.

8. Wang, X., et al., Genome-wide association study identifies QTL for thousand grain weight in winter wheat under normal- and late-sown stressed environments. Theoretical and Applied Genetics, 2021. 134(1): p. 143–157.

9. Ullah, A., et al., Heat stress effects on the reproductive physiology and yield of wheat. Journal of Agronomy and Crop Science, 2022. 208(1): p. 1–17.

10. Barkley, A., et al., The impact of climate, disease, and wheat breeding on wheat variety yields in Kansas, 1985–2011. Bulletin, 2013. 665: p. 1–31.

11. Bergkamp, B., et al., Prominent winter wheat varieties response to post-flowering heat stress under controlled chambers and field based heat tents. Field Crops Research, 2018. 222: p. 143–152.

12. Hütsch, B.W., D. Jahn, and S. Schubert, Grain yield of wheat (Triticum aestivum L.) under long-term heat stress is sink-limited with stronger inhibition of kernel setting than grain filling. Journal of Agronomy and Crop Science, 2019. 205(1): p. 22–32.

13. Wang, X., et al., Physiological and proteome studies of responses to heat stress during grain filling in contrasting wheat cultivars. Plant Science, 2015. 230: p. 33–50.

14. Porter, J.R. and M. Gawith, Temperatures and the growth and development of wheat: a review. European Journal of Agronomy, 1999. 10(1): p. 23–36.

15. Porter, J.R., Rising temperatures are likely to reduce crop yields. Nature, 2005. 436(7048): p. 174–174.

16. Kirby, E.J.M., Ear development in spring wheat. The Journal of Agricultural Science, 1974. 82(3): p. 437–447.

17. Prasad, P.V.V. and M. Djanaguiraman, Response of floret fertility and individual grain weight of wheat to high temperature stress: sensitive stages and thresholds for temperature and duration. 2014(1445-4416 (Electronic)).

18. Dias, A.S. and F.C. Lidon, Evaluation of Grain Filling Rate and Duration in Bread and Durum Wheat, under Heat Stress after Anthesis. Journal of Agronomy and Crop Science, 2009. 195(2): p. 137–147.

19. Wardlaw, I.F., I. Sofield, and P. Cartwright, Factors Limiting the Rate of Dry Matter Accumulation in the Grain of Wheat Grown at High Temperature. Australian Journal of Plant Physiology, 1980. 7: p. 387–400.

20. Wardlaw, I.F. and L. Moncur, The Response of Wheat to High Temperature Following Anthesis. I. The Rate and Duration of Kernel Filling. Australian Journal of Plant Physiology, 1995. 22: p. 391–397.

21. Sharrow, S.H., A Simple Disc Meter for Measurement of Pasture Height and Forage Bulk. Journal of Range Management, 1984. 37(1): p. 94–95.

22. R development core team, R., R development core team. RA Lang Environ Stat Comput, 2013. 55: p. 275–286.

23. Mendiburu, F. and R. Simon, agricolae. 2009.

24. Kuznetsova, A., P.B. Brockhoff, and R.H.B. Christensen, lmerTest Package: Tests in Linear Mixed Effects Models. Journal of Statistical Software, 2017. 82(13): p. 1 – 26.

25. Simko, T.W.a.V. R package ‘corrplot’: Visualization of a Correlation Matrix. 2021; Available from: https://github.com/taiyun/corrplot.

26. Ginestet, C., ggplot2: Elegant Graphics for Data Analysis. Journal of the Royal Statistical Society Series A: Statistics in Society, 2011. 174(1): p. 245–246.

27. Bassu, S., et al., Optimising sowing date of durum wheat in a variable Mediterranean environment. Field Crops Research, 2009. 111(1): p. 109–118.

28. Padovan, G., et al., Understanding effects of genotype × environment × sowing window interactions for durum wheat in the Mediterranean basin. Field Crops Research, 2020. 259: p. 107969.

29. Ul-Allah, S., et al., Phenotypic characterization of wheat germplasm for heritability and dissection of association among post anthesis traits under variable sowing dates. Journal of King Saud University - Science, 2023. 35(3): p. 102578.

30. Hunt, J.R., et al., Early sowing systems can boost Australian wheat yields despite recent climate change. Nature Climate Change, 2019. 9(3): p. 244–247.

31. Wajid, A., et al. Crop Models: Important Tools in Decision Support System to Manage Wheat Production under Vulnerable Environments. Agriculture, 2021. 11, DOI: 10.3390/agriculture11111166.

32. Prieto, P., et al., Dynamics of floret initiation/death determining spike fertility in wheat as affected by Ppd genes under field conditions. Journal of Experimental Botany, 2018. 69(1460-2431 (Electronic)): p. 2633–2645.

33. Youssefian, S., E.J.M. Kirby, and M.D. Gale, Pleiotropic effects of the GA-insensitive Rht dwarfing genes in wheat. 2. Effects on leaf, stem, ear and floret growth. Field Crops Research, 1992. 28(3): p. 191–210.

34. Reynolds, M., et al., Achieving yield gains in wheat. Plant, Cell and Environment, 2012. 35,(1365-3040 (Electronic)): p. 1799–1823.

35. Griffiths, S., et al., Genetic dissection of grain size and grain number trade-offs in CIMMYT wheat germplasm. Plose One, 2015. 16(1932-6203 (Electronic)): p. 18.

36. Zhang, B., et al., Estimation of grain filling rate and thousand-grain weight of winter wheat (Triticum aestivum L.) using UAV-based multispectral images. European Journal of Agronomy, 2024. 159: p. 127258.

37. Elbashier, E., Idris, S., Tahir, I., Elhanafi, S., Saad, A., Mustafa, H., Idris, A. and Tadesse, W., Genome Wide Association Study of Yield and Yield-Related Traits in Elite Spring Bread Wheat Genotypes Grown under High Temperature Environment in Sudan. American Journal of Plant Sciences, 2023. 14: p. 18.

38. Sareen, S., et al. Resilience to Terminal Drought, Heat, and Their Combination Stress in Wheat Genotypes. Agronomy, 2023. 13, DOI: 10.3390/agronomy13030891.

39. Shaalan, A., M.A. Attia, and M.A. Hassaan, Response of Some Wheat Cultivars to Sowing Dates and Biofertilizers under North West Coast of Egypt. Egyptian Journal of Agronomy, 2019. 41(3): p. 313–324.

40. Basheir, S.M., et al. Identification of Wheat Germplasm Resistance to Late Sowing. Agronomy, 2023. 13, DOI: 10.3390/agronomy13041010.

41. Kumar, S., et al., Capturing agro-morphological variability for tolerance to terminal heat and combined heat–drought stress in landraces and elite cultivar collection of wheat. Frontiers in Plant Science, 2023. Volume 14 -2023.

42. Rehman, H.U., et al. Evaluation of Physiological and Morphological Traits for Improving Spring Wheat Adaptation to Terminal Heat Stress. Plants, 2021. 10, DOI: 10.3390/plants10030455.

43. Gupta, V., et al., AMMI and GGE biplot analysis of yield under terminal heat tolerance in wheat. (1573-4978 (Electronic)).

44. Devi, K., et al., Identification of Wheat Genotypes Resilient to Terminal Heat Stress Using GGE Biplot Analysis. Journal of Soil Science and Plant Nutrition, 2022. 22(3): p. 3386–3398.

45. Vicentin, L., J. Canales, and D.F. Calderini, The trade-off between grain weight and grain number in wheat is explained by the overlapping of the key phases determining these major yield components. Frontiers in Plant Science, 2024. Volume 15 -2024.

46. Kim, J., R. Savin, and G.A. Slafer, Quantifying pre- and post-anthesis heat waves on grain number and grain weight of contrasting wheat cultivars. Field Crops Research, 2024. 307: p. 109264.

47. Qaseem, M.F., R. Qureshi, and H. Shaheen, Effects of Pre-Anthesis Drought, Heat and Their Combination on the Growth, Yield and Physiology of diverse Wheat (Triticum aestivum L.) Genotypes Varying in Sensitivity to Heat and drought stress. Scientific Reports, 2019. 9(1): p. 6955.

